# Systematic comparison and rational design of theophylline riboswitches for efficient gene repression

**DOI:** 10.1101/2022.07.17.500366

**Authors:** Xun Wang, Can Fang, Yifei Wang, Xinyu Shi, Fan Yu, Jin Xiong, Shan-Ho Chou, Jin He

**Affiliations:** State Key Laboratory of Agricultural Microbiology, College of Life Science and Technology, Huazhong Agricultural University, Wuhan, Hubei 430070, P. R. China

## Abstract

Riboswitches are promising regulatory tools in synthetic biology. To date, 25 theophylline riboswitches have been developed for gene expression regulation in bacteria. However, no one has systematically evaluated their regulatory effects. To facilitate rational selection of theophylline riboswitches, we examined 25 theophylline riboswitches in *Escherichia coli* and surprised to find that none of the five repressive riboswitches were more than 2-fold effective. To solve this problem, we rationally designed a transcriptional repressive riboswitch and demonstrated its effect not only in various bacterial strains but also in different growth media or different temperatures. By introducing two copies of theophylline riboswitches and a RepA protein degradation tag coding sequence at the 5’-end of a reporter gene, we successfully constructed a dual gene expression regulatory system with up to 150-fold potency, namely the R2-RepA system. R2-RepA system is only 218 bp in length, expression of any protein could be repressed efficiently by simply inserting this system upstream of the target protein-coding sequence. This study represented a crucial step toward harnessing theophylline riboswitches and expanding the synthetic biology toolbox.

## INTRODUCTION

Riboswitches are common gene regulatory elements typically located in the 5’-untranslated region (UTR) of mRNAs that alter gene expression in response to small molecule ligands (Winkler *et al*, 2002). It comprises two parts, an aptamer domain that binds ligand and an expression platform that regulates the expression of downstream genes (Breaker, 2012). Due to their simplicity, specificity, stability, modular design, and ease of implementation, riboswitches provide a promising platform for gene regulation.

To meet the growing demand, more than 60 artificial riboswitches that respond to non-metabolite ligands, including theophylline, tetracycline, naringenin, caprolactam and dopamine, have been constructed (Borujeni *et al*, 2016; Harbaugh *et al*, 2022; Jang *et al*, 2019; Jang *et al*, 2017; Suess *et al*, 2004; Suess *et al*, 2003). Of these, theophylline riboswitch is the most studied (Wrist *et al*, 2020). In 2004, Suess et al. engineered the first theophylline riboswitch by combining theophylline aptamer with an expression platform, resulting in a functional translational ON (TL-ON) riboswitch (Suess *et al.,* 2004). Shortly thereafter, Desai and Gallivan constructed another TL-ON theophylline riboswitch (Desai & Gallivan, 2004). Later, other researchers developed theophylline riboswitches with different regulatory mechanisms. For examples, Ogawa et al. fused theophylline aptamer to a hammerhead ribozyme to generate a ribozyme ON (RZ-ON) riboswitch (Ogawa & Maeda, 2008), and Fowler et al. selected the first transcriptional ON (TC-ON) theophylline riboswitch by fluorescence-activated cell sorting (FACS) technique (Fowler *et al*, 2008). Topp and Gallivan further constructed a translational OFF (TL-OFF) theophylline riboswitch by inserting a riboswitch coding sequence within the translated region of a gene (Topp & Gallivan, 2008). On top of that, Ceres et al. rational designed three chimeric riboswitches, each containing the same theophylline aptamer domain, with three independent expression platforms from *metE, yitJ* and *lysC* riboswitches to generate three functional transcriptional OFF (TC-OFF) riboswitches (Ceres *et al*, 2013a). Meanwhile, various research teams have extensively screened and optimized the theophylline riboswitch library to improve their regulatory ability. In order to accurately characterize the regulatory ability of riboswitches, researchers introduced activation/inhibition ratio, which refers to the quantitative relationship between small molecule inducer concentration and biosensor output signal (Snoek *et al*, 2020). The activation/repression ratio is calculated as the fold change between the maximum and minimum values of biosensor output signal. For example, Topp et al. screened and constructed six TL-ON theophylline riboswitches termed A-E and E*, which enable inducible gene expression in eight different bacterial species with activation ranging from 5 to 150-fold (Topp *et al*, 2011). Cui et al. modified the ribosome binding site (RBS) in riboswitch E to generate riboswitch E1 that results in an activation fold of 6.8 in *Bacillus subtilis* (Cui *et al*, 2016). After that, Canadas et al. generated four riboswitches based on the sequence of riboswitch E*, providing effective activation in *Clostridium* (Canadas *et al*, 2019). Details of the above riboswitches are shown in Table 1.

**Table 1.**
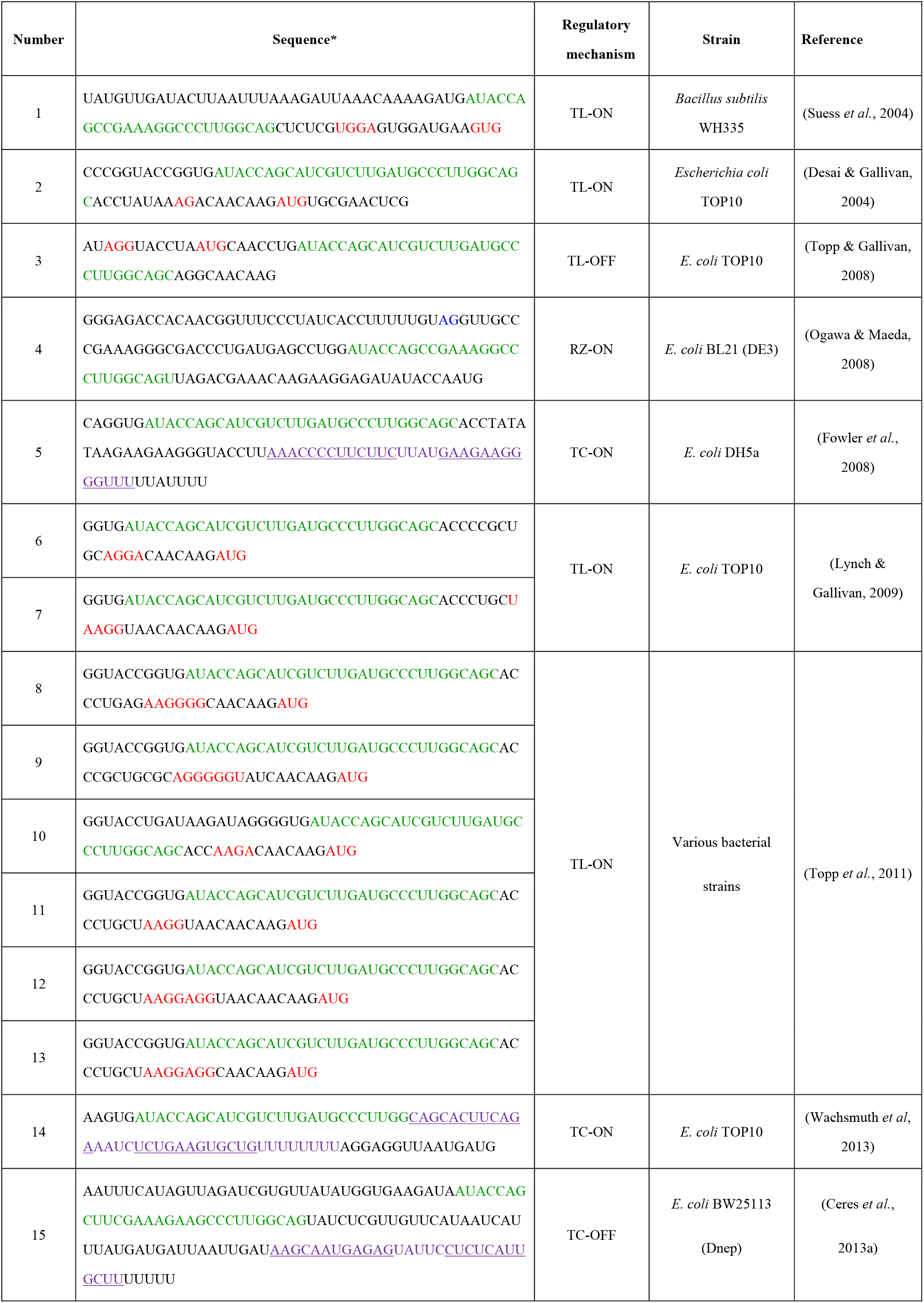
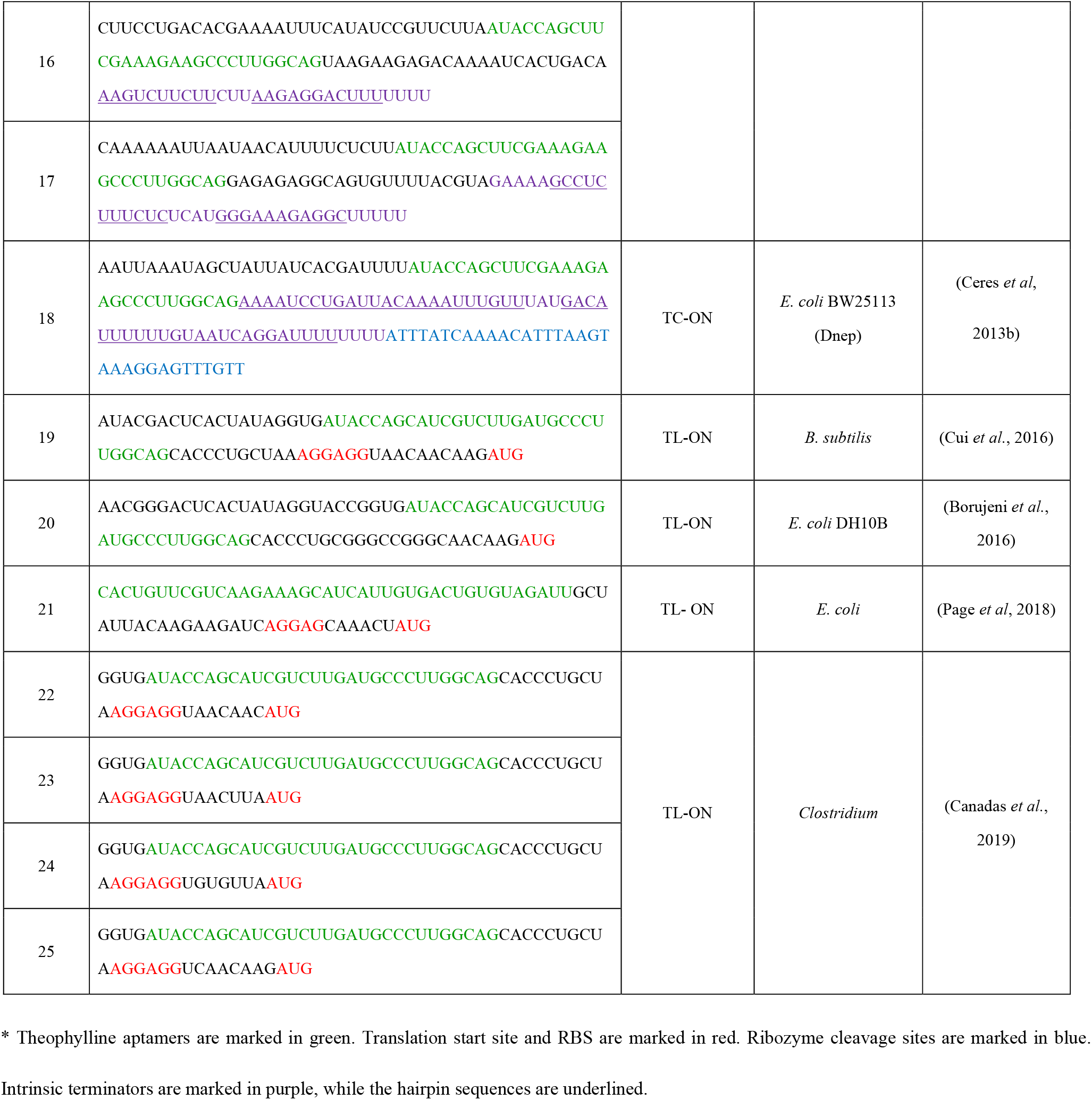
A list of theophylline riboswitches used in bacteria published to date.

From the above, it can be concluded that theophylline riboswitches have been developed for five distinct regulatory mechanisms that can mediate two different regulatory effects: activation or repression of gene expression. While many studies have assessed the activation/repression ratios of these riboswitches under various conditions, systematic comparisons of their regulatory functions under the same experimental condition are lacking. To facilitate a more rational selection of theophylline riboswitches, we focused on two important parameters, the activation/repression ratio and the basal expression level. The latter is the output signal of the biosensor in the absence of the inducer (Rogers *et al*, 2015). We examined 25 theophylline riboswitches commonly used in bacterial cells, including 17 TL-ON, 3 TC-ON, 1 RZ-ON, 1 TL-OFF, and 3 TC-OFF riboswitches. We investigated these two parameters in *Escherichia coli* MG1655 strain and found that they were highly variable. We also compared the data in different *E. coli* strains, growth media, and temperatures, and found, notably, that the regulatory effects of riboswitches were unsatisfactory except for TL-ON riboswitches. To obtain an efficient repressive riboswitch, we rationally designed and constructed a novel TC-OFF theophylline riboswitch. To further enhance the effect, we employed two strategies: first, we connected riboswitches in tandem to increase their activation/repression ratios; second, we exploited a protein degradation tag to shorten the half-life of the proteins, thereby achieving extremely low protein leaky expression. We also constructed a mathematical model to predict the systemic repression effects at different theophylline concentrations. This work thus provided a new biological cassette for bottom-up design of genetic circuits that shall greatly facilitate rational engineering of gene expression in synthetic and systems biology.

## MATERIAL AND METHODS

### Plasmid construction

In *E. coli* and *S*. Typhimurium, reporter plasmids were constructed using pBRplac as the parent plasmid (Beisel & Storz, 2011; Guillier & Gottesman, 2006). In *M. smegmatis, B. thuringiensis* and *B. subtilis,* plasmids pMV261 (Ali *et al*, 2017; Li *et al*, 2022), pRP0122 (Zhou *et al*, 2016) and pHT43 (Rafique *et al*, 2021) were used respectively. Except for pHT43, the transcription of *turborfp* along with its 5’UTR regulatory elements was carried out under strong constitutive promoters. In pHT43, gene expression was controlled by the strong IPTG-inducible Pgrac promoter. The primers used for plasmid construction were synthesized by Tianyi Huiyuan (Wuhan, Hubei, China). All plasmids generated in this study were assembled using the Hieff Clone^®^Plus Multi One Step Cloning Kit (YEASEN, Shanghai, China), and confirmed *via* sequencing (Quintarabio, Wuhan, Hubei, China). The constructed plasmids were transformed into corresponding strains via calcium chloride (CaCl_2_) method to produce corresponding derivative strains (Supplementary Table S1). The plasmids and primers used in this study were listed in Appendix Table S2 and S3.

### Bacteria and culture conditions

*E. coli* DH5α was used for all cloning experiments. If not indicated otherwise, *E. coli* MG1655 was transformed with the resulting plasmids for fluorescence measurement. *E. coli* NST74 (Tribe, 1987), BL21, HB101, JM101, BW25113 and Top10, *Salmonella enterica* serovar Typhimurium SL1344 (Richardson *et al*, 2011), *Mycobacteria smegmatis* MC^2^155 (Li *et al*, 2017), *Bacillus thuringiensis* BMB171 (He *et al*, 2010; Wang *et al*, 2019) and *Bacillus subtilis* 168 were employed as hosts for riboswitch performance tests (Appendix Table S1). *E. coli*, *S.* Typhimurium and *B. thuringiensis* were grown in lysogeny broth (LB) medium (tryptone 10 g/L; yeast extract 5 g/L; NaCl 10 g/L), and *M. smegmatis* in 7H9 media (7H9 Broth 4.9 g/L; 0.2% glycerol; Tween 20 0.05%). For *B. subtilis,* 2×yeast extract tryptone (2×YT) medium (tryptone 16 g/L; yeast extract 10 g/L; NaCl 5 g/L) were used for cultivation. When necessary, ampicillin, kanamycin and spectinomycin was added to the culture at the final concentrations of 100 μg/mL, 50 μg/mL, and 100 μg/mL, respectively. If not otherwise indicated, strains were grown in 250 mL shake flasks (50 mL medium per flask) on a rotary shaker at 200 rpm at 37 °C (Ruihua, Wuhan, Hubei, China).

For fluorescence measurement, individual colonies were picked and grown overnight in 5 mL LB media with 100 μg/mL ampicillin. This culture was diluted 100-fold to inoculate 50 mL of fresh media and allowed to grow to early exponential phase (2 hours after inoculation, an OD_600_ approximately 0.5), at which point theophylline was added to the media at the concentrations indicated. Cells were allowed to grow at 37 °C for different times to measure their fluorescence intensity as indicated in the figure legend.

### RNA extraction, cDNA synthesis, and RT-qPCR

For RT-qPCR experiments, 2 mL samples from *E. coli* strains were collected. Total RNA was extracted, and RT-qPCR was conducted essentially as previously described (Wang *et al*, 2014), with modifications as indicated in the figure legends. Results for each strain were normalized to those of the *rrsB* gene coding 16S rRNA. For data analysis, technical and biological triplet data were obtained. Data were subjected to one-way analysis of variance (ANOVA) using the Bonferroni test, n = 3.

### Fluorescence measurement

Samples were taken in triplicate (1 mL each sample), and OD_600_ was measured using 500 μL of culture. Another 500 μL of the culture was taken and diluted to appropriate concentration with LB media to measure its fluorescence intensity in a 96 well microplate (Sangon Biotech, Shanghai, China). Fluorescent measurements were carried out at an excitation wavelength of 553 nm, and the emission fluorescence was taken at 593 nm. All fluorescence intensity results were normalized by respective cell growth (OD_600_) and background elimination. All experimental results were obtained with three biological replicates. Data were subjected to one-way analysis of variance (ANOVA) using the Bonferroni test, n = 3.

### Mathematical model

The model was generated in MATLAB R2019b. The equations used were based on the law of mass action that describes the biomolecular interactions. A detailed derivation and description of the model was provided in the results section and in the Appendix file (Koch, 1956; Quand *et al*, 2013).

## RESULTS

### Systematic evaluation of theophylline riboswitches

To test the regulatory role of each riboswitch, we first determined the applied host strain, culture conditions, and appropriate inducer concentrations. We selected the most representative “wild type” *E. coli* strain MG1655, and the corresponding LB medium, incubated it at 37°C. We then examined the inhibitory effect of theophylline on the growth of MG1655 and found that the growth inhibition in LB medium was negligible when theophylline concentration was less than 2 mM (Appendix Fig S1). Therefore, 2 mM theophylline was used in all following experiments unless otherwise indicated. We compiled the detailed sequences of 25 theophylline riboswitches commonly used in bacteria by reviewing the literature and classified them into 17 TL-ON, 3 TC-ON, 1 RZ-ON, 1 TL-OFF, and 3 TC-OFF riboswitches according to their regulatory mechanisms and effects (Tables 1). We then separately inserted coding sequences of different theophylline riboswitches upstream of the reporter gene *turborfp* controlled by a constitutive promoter (Fig 1A) to construct 25 different plasmids containing theophylline riboswitches (Appendix Table S2). We also constructed the pWA143 plasmid containing the gene circuit but without the riboswitch coding sequence as a negative control (Fig 1A). We then transformed these plasmids individually into MG1655 to test the function of different theophylline riboswitches.

**Figure 1.**
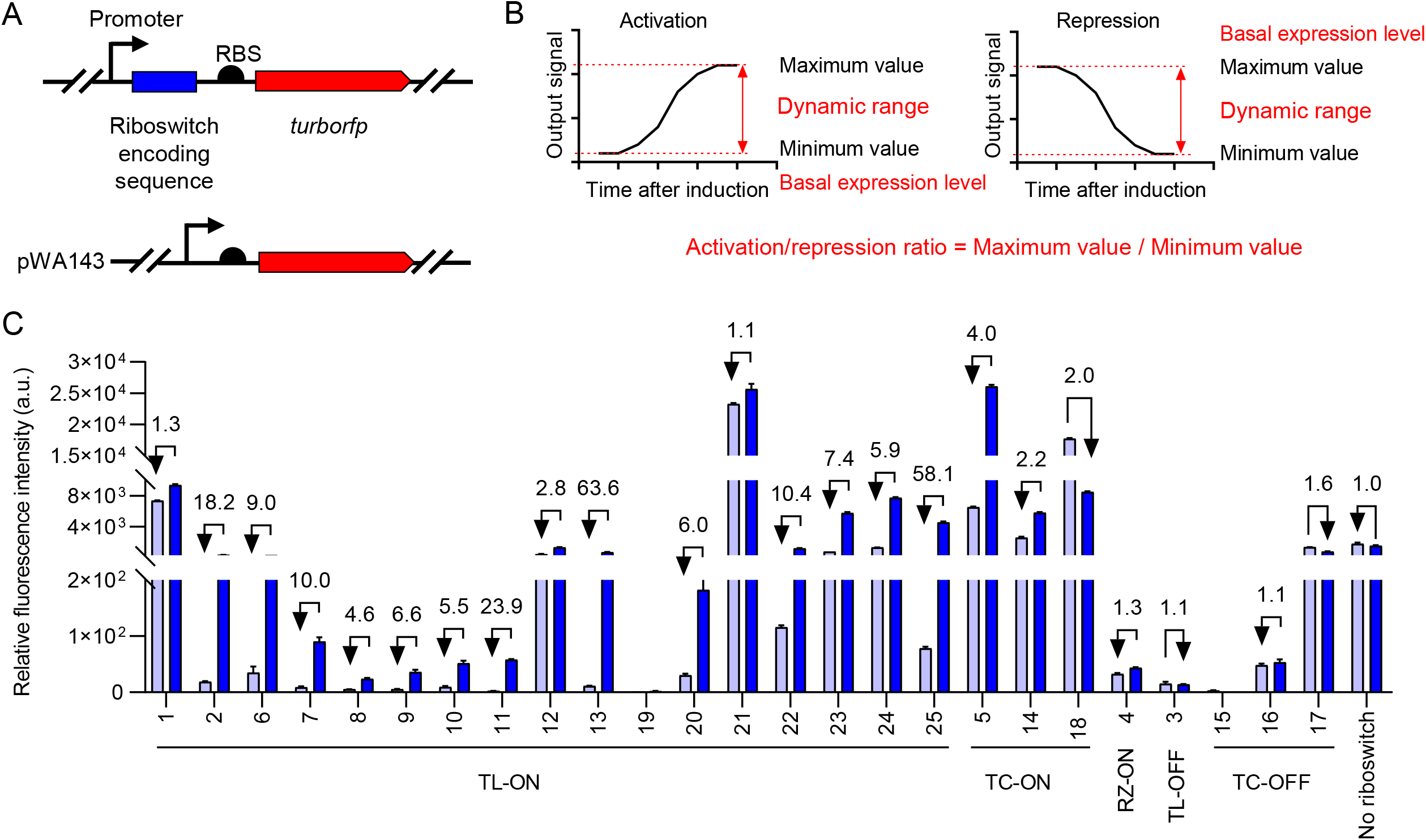
Evaluation of gene expression regulation efficiencies of various theophylline riboswitches. A Designed plasmids containing gene circuits to assess the repression profile of theophylline riboswitches. The *turborfp* gene (shown as red arrow) is controlled by a constitutive promoter (shown as black arrow). Also shown are the theophylline riboswitch coding sequence (blue box) and RBS coding sequence (black semicircle). Control plasmid pWA143, including the gene circuit without the riboswitch coding sequence. B Definition of activation/repression ratio, basal expression level, and dynamic range. The activation/repression ratio is calculated by dividing the maximum value with the minimum value. Basal expression level refers to the output signal prior to addition of the inducer. In the case of activation, the minimum value is equal to the basal expression level; in the case of repression, the maximum value is equal to the basal expression level. Dynamic range refers to the maximum and minimum values of the interval. C Relative expression levels of TurboRFP fluorescence intensity measured in the absence (light blue) and presence (dark blue) of 2 mM theophylline. The numbers above the column represent activation/repression ratios. Data represent mean ± SD of 3 biological replicates.

To evaluate these riboswitches, we examined their performance on activation/repression ratios and basal expression levels. Activation/repression ratio was calculated by dividing the TurboRFP fluorescence intensity at 2 mM theophylline by the fluorescence intensity at 0 mM theophylline. Basal expression levels refer to TurboRFP fluorescence intensity at 0 mM theophylline. These two parameters together constitute the dynamic range of the riboswitch, that is, the maximum and minimum values that it can regulate (Fig 1B).

First, we focused on the activation/repression ratios of these riboswitches (Fig 1C). Most 17 TL-ON riboswitches showed 2.2- to 63-fold activation; unfortunately, No. 1 and No. 21 exhibited less than 2-fold activation, and No. 19 had almost no fluorescent signal. Of the 3 TC-ON riboswitches, No. 5 and No. 14 promoted more than 2-fold expression of TurboRFP in the presence of theophylline, while No. 18 showed the opposite effect under our experimental condition; it did not activate, but inhibited gene expression. For No. 4 riboswitch of RZ-ON, the data showed that it barely worked. It can be seen that of the above 21 ON-switches, including 17 TL-ON, 3 TC-ON and 1 RZ-ON, No. 13 exhibited the highest activation ratio of 63.6-fold. Of the 4 OFF-switches, No. 3 riboswitch was reported to repress translation, but was only 1.1-fold effective when grown in the presence of theophylline. Three TC-OFF riboswitches (No. 15, No. 16, and No. 17) were also tested. Only No. 17 showed a 1.6-fold difference in repressing gene expression, and the other two had almost unchanged TurboRFP fluorescence intensities under theophylline induction.

Next, we compared basal expression levels and found that they varied widely (Fig 1C). Among the “ON” riboswitches, the relative basal levels of TurboRFP fluorescence expression ranged from as low as 2.5 arbitrary units (a.u.) (No. 11) to as high as 23,000 a.u. (No. 21), meaning that they differ by a factor of almost 10,000. In the “OFF” riboswitches, there is also a large difference in the basal expression levels of TurboRFP. For example, the highest (No. 17) and the lowest (No. 3) were 1,300 a.u. and 15 a.u., respectively. Given the limited activation/repression ratio of most riboswitches, these differences cannot be ignored. For example, if two riboswitches both activated gene expression up to 10-fold, and if one has a basal expression level of 100 a.u. and the other of 1000 a.u., then they regulated with a dynamic range of 100-1,000 a.u., and 1,000-10,000 a.u., respectively, which could lead to a large difference in gene expression. That said, when we focused on the activation/repression ratios of riboswitches, we should also pay attention to their basal expression levels.

Finally, we tested the performance of the aforementioned 25 riboswitches under different conditions, including different *E. coli* strains, different temperatures, and different media. Although the degrees of activation/repression have changed, it is worth noting that for most riboswitches, dynamic range didn’t change much. For example, for TL-ON riboswitch No. 6, in strain JM101, the activation ratio in LB medium at 37 °C was only 1.1, but in strain DH5a, it reached 9.6 in LB medium at 25 °C. Although far different, the relative TurboRFP fluorescence intensities were still in the dynamic range of 100-300 a.u. This was also true for other riboswitches, such as TL-ON riboswitch No. 21, where the dynamic range was consistently within 20,000-40,000 a.u., regardless of the conditions. Therefore, testing riboswitches under various conditions allowed us to more accurately assess their functions.

Notably, TL-ON riboswitches No. 13 showed superior results, with activation ratio exceeding 12 under all tested conditions. Its basal expression ratios fluctuated in the range of 3-76 a.u. (Appendix Fig S2). Therefore, this riboswitch was the best player of all ON-riboswitches. On the contrary, all OFF-riboswitches performed poorly, with repression ratio less than 2-fold. For TC-OFF riboswitch No. 17, previous results showed that its repression ratio was 1.6 at 2 mM theophylline than at 0 mM theophylline, and was ineffective at 25°C and in SOC medium.

From the above data, we could conclude that among the 5 regulatory mechanisms, TL-ON riboswitches were the best optimized, especially riboswitches No. 13, because it showed the highest activation ratio and the lowest basal expression level. However, none of the “OFF” riboswitches showed over 2-fold repression ratio, so we needed to redesign and reconstruct a repressive riboswitch.

### Rational design of TC-OFF theophylline riboswitch

Among the riboswitches for transcriptional regulation, there are two types of regulation based on intrinsic terminators and those based on Rho-dependent terminators (Proshkin *et al*, 2014; Wang *et al.*, 2019). Intrinsic terminators are sequences in the non-template DNA strand that, when transcribed into RNA, forms a GC-rich hairpin structure followed by a U-rich tract in the RNA:DNA hybrid (Rosenberg & Court, 1979). It leads to dissociation of elongation complex without the assistance of auxiliary transcription regulators. Compared to translational and ribozyme-based regulation, transcriptional control by intrinsic termination is a more conserved, relatively simple, and efficient regulatory mechanism (Mitra *et al*, 2009). Considering the advantages of intrinsic terminator-based transcriptional control, we sought to develop a TC-OFF theophylline riboswitch. Riboswitch B (No. 9 in this work) was previously reported to activate protein translation (Topp *et al.*, 2011). In the absence of theophylline, riboswitch B folds into an OFF state in which the RBS was sequestered in the secondary structure. When theophylline binds to the aptamer, RBS becomes accessible to 16S rRNA of the ribosome to initiate translation (Topp *et al.,* 2011). Interestingly, we found that the RBS sequence “AGGGGGU” is rich in G, which represents exactly half of the intrinsic transcription terminator hairpin sequence. We speculated that if the downstream sequence of RBS was replaced by another half of the terminator, a complete intrinsic transcription terminator could be constructed. Therefore, we changed the sequence “CAAGAUG” to “CCCCCUU” and added another 7 U residues (UUUUUUU) downstream of it. The terminator is thus composed of a 7 bp hairpin stem, a 4 bp loop, and a stretch of 8 U residues (Fig 2A). We named this new riboswitch R1 and expected it to transcriptionally repress expression. Since the translation initiation codon AUG in riboswitch B was deleted in the new construct, we added an RBS and an AUG codon downstream of the riboswitch to initiate TurboRFP translation. The corresponding plasmid was named pWA131 (Appendix Fig S3, Table S2).

**Figure 2.**
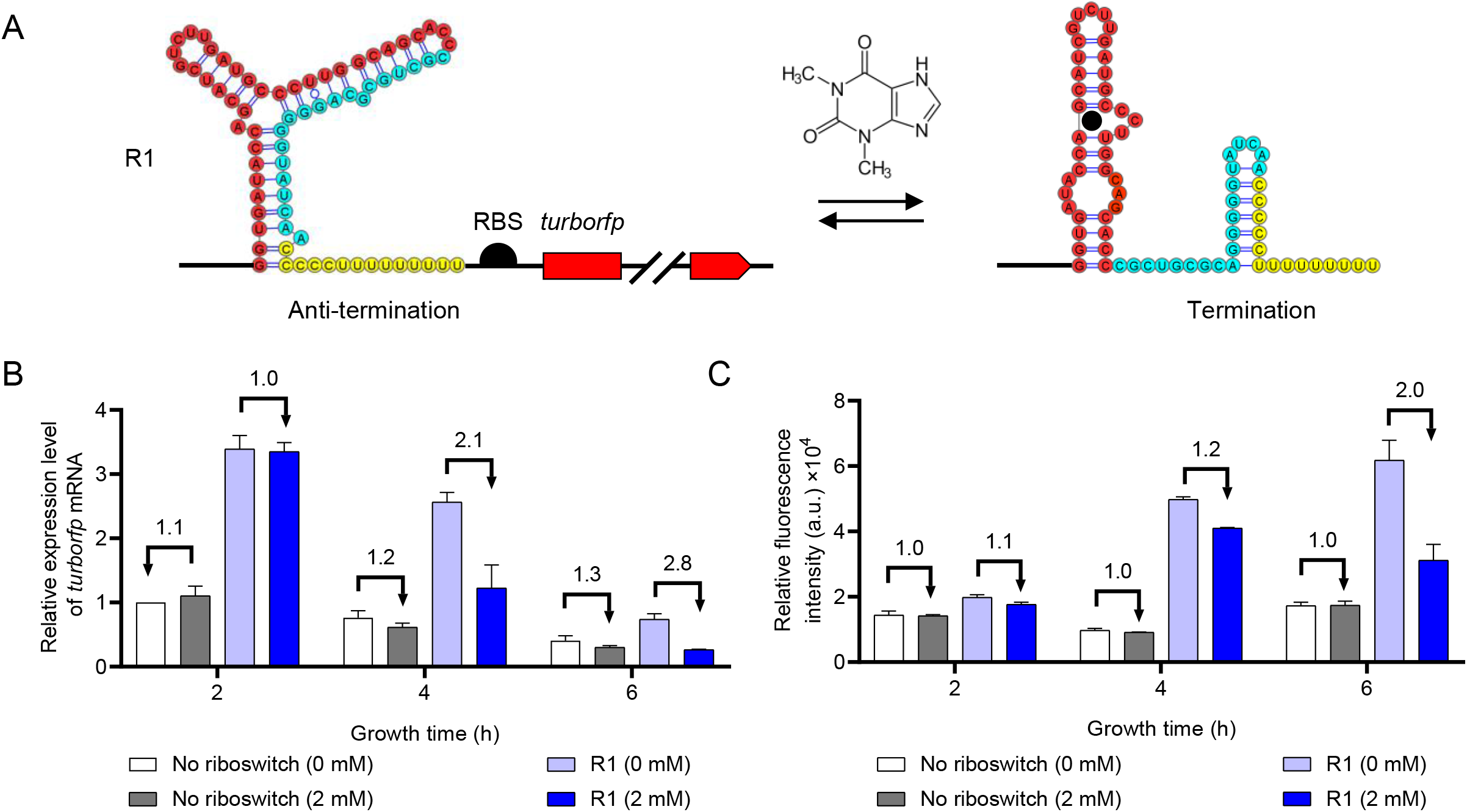
Evaluation of gene expression regulation efficiencies of rationally designed TC-OFF theophylline riboswitches. A Design strategy for theophylline-dependent riboswitch R1 to control transcription. Theophylline aptamer (red) was fused to an intrinsic transcription terminator (cyan and yellow). Sequences modified from the previous riboswitch B were marked in yellow. The RBS sequence (black semicircle) and the open reading frame of the reporter gene *turborfp* are located downstream of this construct. In the absence of theophylline, intrinsic terminator formation is inhibited, resulting in transcription readthrough and *turborfp* expression. Upon binding of theophylline (black solid circle), an intrinsic terminator is formed and transcription is prematurely stopped, resulting in repression of *turborfp* expression. B Relative expression levels of *turboRFP* mRNA measured in the absence (white and light blue) and presence (gray and dark blue) of 2 mM theophylline. C Relative TurboRFP fluorescence intensities measured in the absence (white and light blue) and presence (gray and dark blue) of 2 mM theophylline. The numbers above the column represent activation/repression ratios. Data represent mean ± SD of 3 biological replicates.

To test whether R1 is functional *in vivo,* MG1655-pWA131 (test strain) or MG1655-pWA143 (control strain) were grown in LB supplemented with 100 μg/mL ampicillin for 2 hours to reach early exponential phase. Theophylline was then added at 2 hours, and *turborfp* mRNA amount or fluorescence density was measured at 2, 4, and 6 hours. The results showed that the expression of *turborfp* mRNA in test strain decreased by 2.1- and 2.8-fold at 4 and 6 hours after the addition of theophylline compared to the case where no theophylline was added. Meanwhile, control strain showed no significant changes in mRNA amount whether theophylline is present or not (Fig 2B). The trends in TurboRFP fluorescence intensities of the test or control strains were consistent with the changes in mRNA amounts. After adding theophylline, the TurboRFP fluorescence intensities decreased by 1.2- and 2.0-fold at 4 and 6 hours in the test train, respectively, compared with the strains without theophylline addition (Fig 2C). Likewise, the TurboRFP fluorescence intensities of the control strain with or without theophylline did not change significantly (Fig 2C). The addition of theophylline thus resulted in a decrease in mRNA and protein levels, demonstrating that our construction of the TC-OFF theophylline riboswitch was indeed working.

### Improvement of riboswitch performance

However, we noticed that the repression ratio of the riboswitch was not large enough. To address this issue, we found previous literature showing that riboswitches in tandem could achieve a greater repression ratio and reduce leaky expression (Sudarsan *et al*, 2006; Zhou *et al.*, 2016). Therefore, we integrated two or three theophylline riboswitches coding sequences linked by a 13 bp linker sequence upstream of *turborfp,* to generate plasmids of pWA140 (R2) and pWA141 (R3), respectively (Fig 3A). Detailed sequences and derivative strains are listed in Appendix Fig S3 and Table S1. At 4 hours, R2 and R3 repressed TurboRFP expression by 1.4- and 1.5-fold in the presence of theophylline. And at 6 hours, these values increased to 3.7- and 2.8-folds, respectively (Fig 3B). From the above data, it can be seen that R2 gave the highest repression ratio among the three at 6 hours. We also found that increasing the number of riboswitches in tandem reduced the basal expression level from 61,830 a.u. (R1) to 39,143 a.u. (R2) to 17,611 a.u. (R3) at 6 hours. After theophylline addition, the corresponding TurboRFP fluorescence intensities also decreased from 31,161 a.u. (R1) to 10,559 a.u. (R2) to 6,395 a.u. (R3).

**Figure 3.**
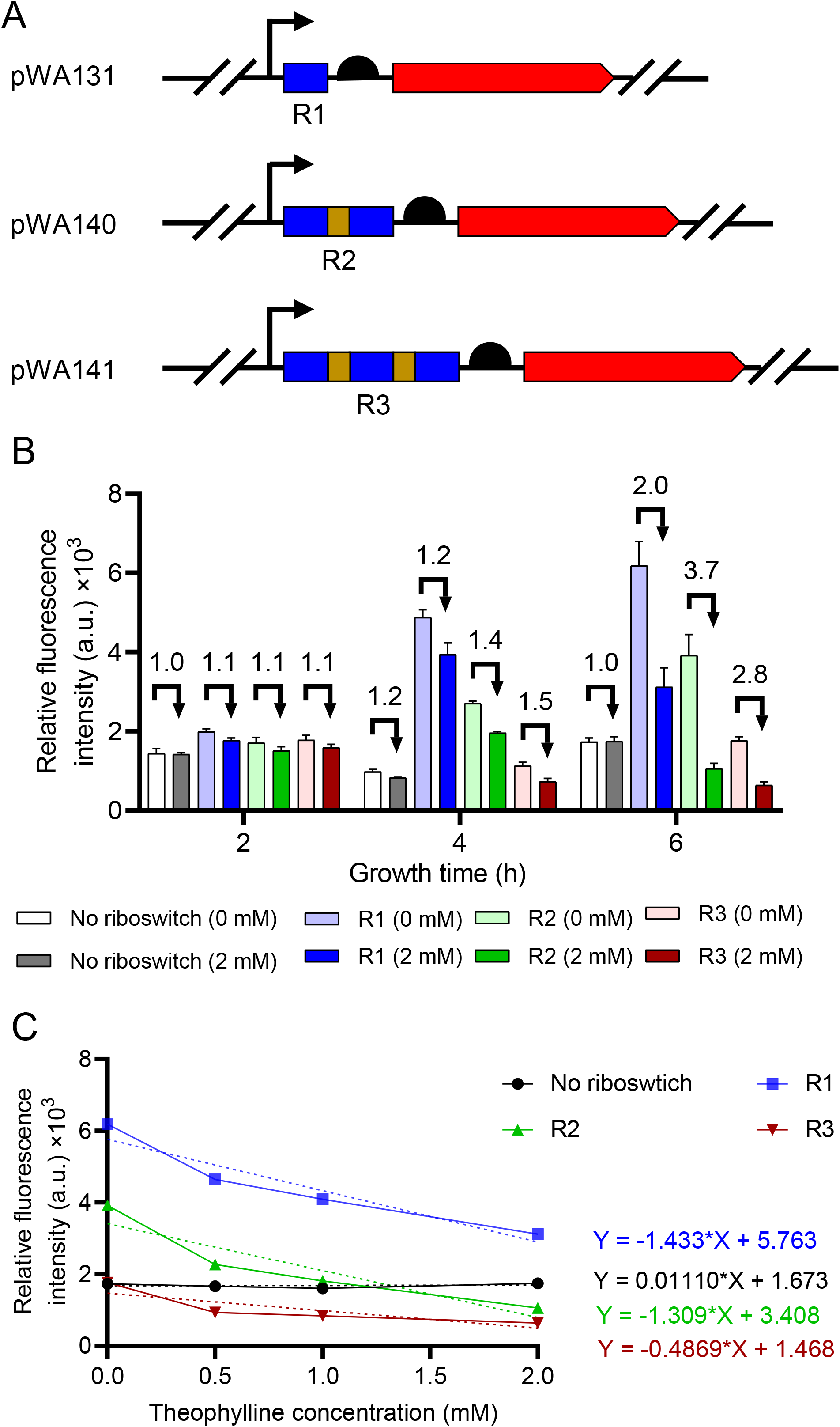
Evaluation of gene expression regulation efficiencies of tandem TC-OFF theophylline riboswitches at different theophylline concentrations. A Schematic of plasmids containing engineered gene circuit controlled by tandem theophylline riboswitches. Promoter (black arrow), riboswitch coding sequences (blue box), linker coding sequences (brown box), RBS coding sequences (black semicircle) and *turboRFP* (red arrow) are indicated. R1 represents one riboswitch, R2 represents two riboswitches in tandem, and R3 represents three riboswitches in tandem. B Relative TurboRFP fluorescence measured in each strain harboring different plasmids with tandem riboswitch (R1, R2 and R3) in the absence and presence of 2 mM theophylline. C Relative TurboRFP fluorescence of each strain grown in LB medium supplemented with different concentrations of theophylline. The numbers above the column represent activation/repression ratios. Data represent mean ± SD of 3 biological replicates.

The response of tandem riboswitches to different concentrations of theophylline was also tested. We measured the fluorescence intensities 6 hours after adding 0, 0.5, 1.0, and 2 mM theophylline. The slopes of the linear regression lines of R1 and R2 were −1.433 and −1.309, respectively, indicating that R1 and R2 showed similar theophylline responses. However, R3 was less sensitive to changes in theophylline concentration, with a slope of the linear regression line of −0.4869 (Fig 3C). Taking into account the repression ratio and basal expression level, we concluded that R2 performed the best among the three.

### Mathematical model of R2-mediated quantifiable repression

To validate our observations and provide predictability, we constructed a mathematic model of the regulatory effect of R2 to different concentrations of theophylline at different times (Fig 4A). As the simulations showed, our model agreed well with the experiment results (R-square was 0.9755). The effect of riboswitch changed greatly when theophylline concentration was below 0.25 mM, and effective repression occurred when theophylline concentration was higher than 0.25 mM. The highest level of repression was reached at 8 hours. Therefore, based on the simulations, we could calculate the fluorescence levels in the theophylline concentration range and time range. Conversely, if the theophylline concentration in the media is unknown, we could deduce the concentration of theophylline in the media from the TurboRFP fluorescence density.

**Figure 4.**
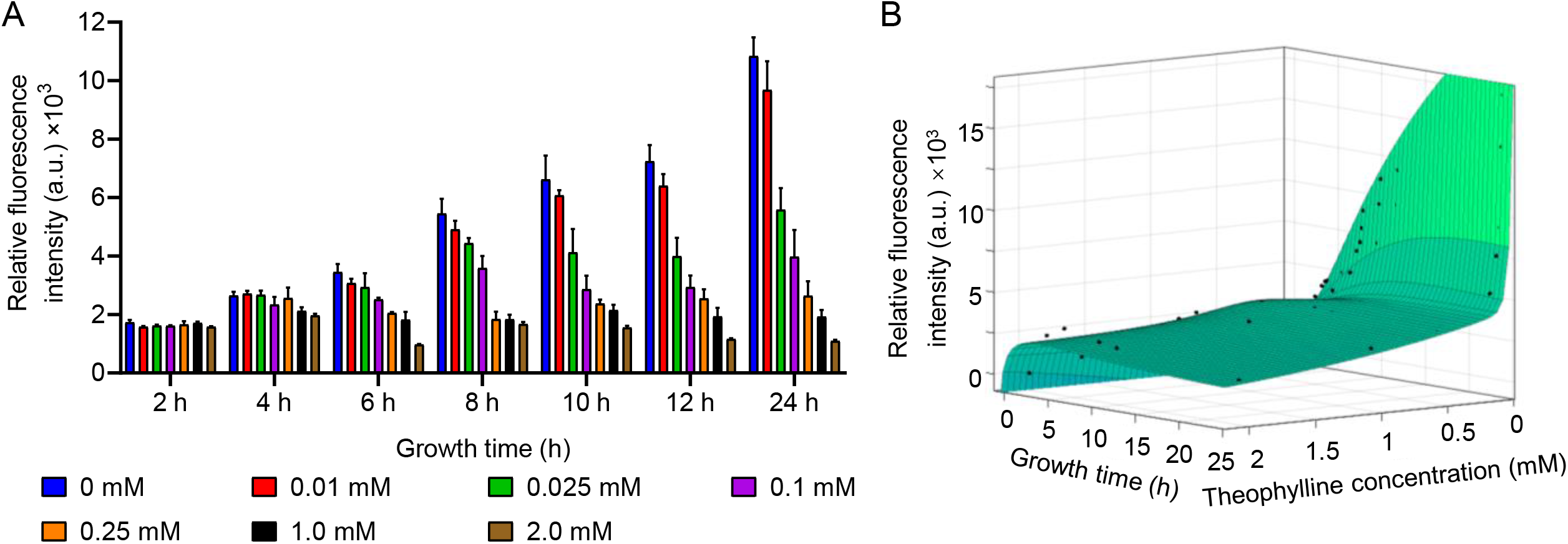
Comparison of experimental measurements and model predictions. A Relative TurboRFP fluorescence measured in strain harboring pWA140 (R2) at various theophylline concentrations from 0 to 2.0 mM over 2 to 24 hours. Data represent mean ± SD of 3 biological replicates. B Mathematic model of R2 based on all data measured at different theophylline concentrations and growth times. Data represent mean ± SD of 3 biological replicates.

### Robustness of R2 under various conditions

To investigate the robustness of R2, we tested it in five different bacteria, including *Proteobacteria E. coli* BL21 and *S. Typhimurium* SL1344, *Actinobacteria M. smegmatis* MC^2^155, *Firmicutes B. subtilis* 168 and *B. thuringiensis* BMB171, which are widely used in genetic engineering. In most strains, R2 provided more than 7-fold repression (Fig 5A). Different *E. coli* stains, including BL21, JM101, HB101, NST74, BW25113, Top10 and DH5α, were also tested. In all strains, the presence of R2 did show a significant repression effect. Among them, JM101 provided the highest repression ratio (Fig 5B).

**Figure 5.**
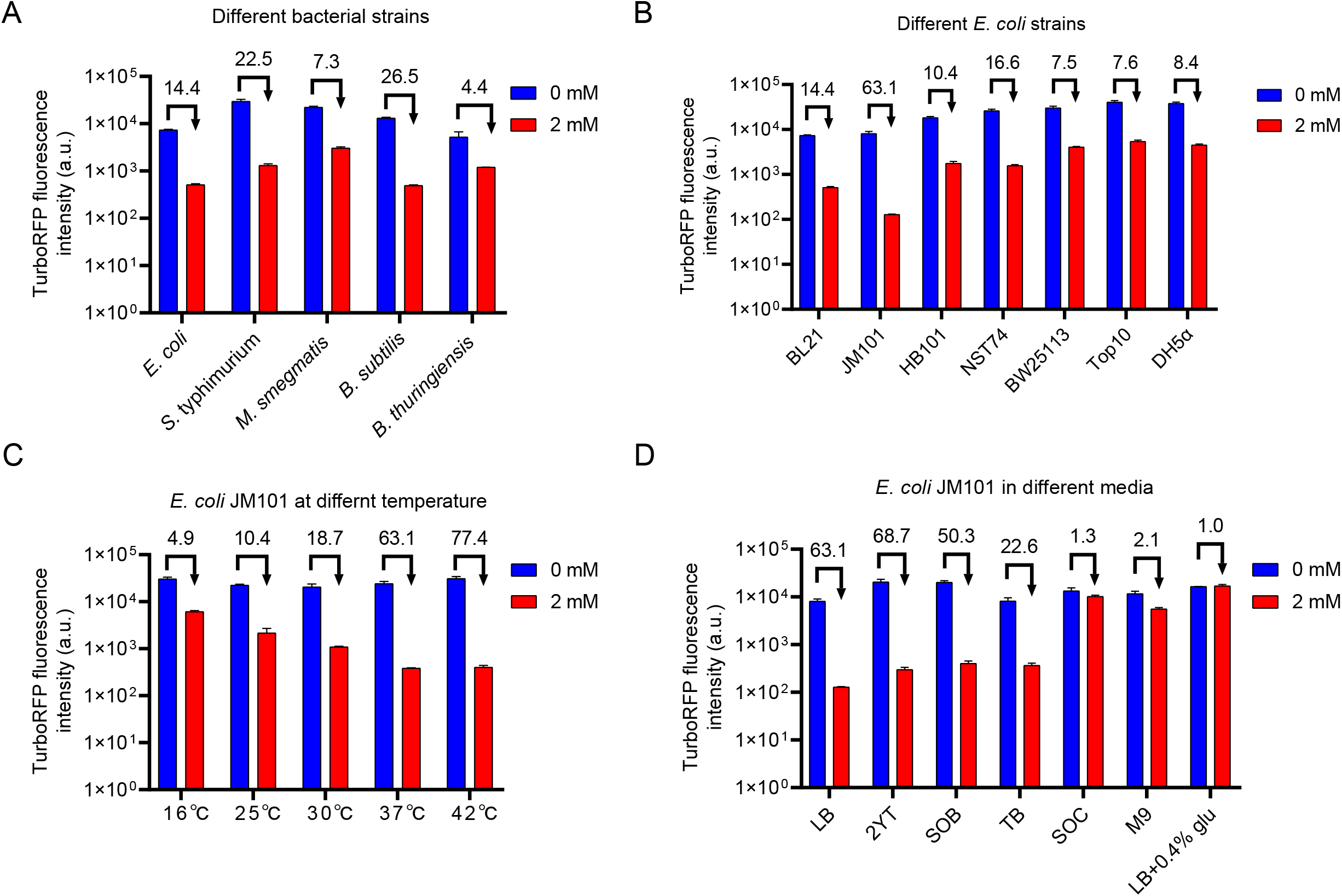
Evaluation of gene expression regulation efficiencies of tandem TC-OFF theophylline riboswitches at different conditions. A Regulation of TurboRFP expression by R2 in different bacterial strains. B Regulation of TurboRFP expression by R2 in different *E. coli* strains. C Regulation of TurboRFP expression by R2 in *E. coli* JM101 strain grown at different temperatures. D Regulation of TurboRFP expression by R2 in *E. coli* JM101 strain grown in different growth media. The numbers above the column represent activation/repression ratios. Data represent mean ± SD of 3 biological replicates.

Since temperature plays an important role in the structure and function of riboswitch RNAs (Fuertig *et al*, 2020), we selected the JM101 strain with the best regulatory effect for testing. R2 in JM101 was tested at 5 different temperatures ranging from 16 °C to 42 °C. Indeed, R2 exhibited significant inhibition at all temperatures tested, but was more potent at 42°C than at 16°C with a factor of 77.4 and 4.9 (Fig 5C). We attributed this phenomenon to a higher mobility of RNA structure at high temperature (Wu *et al*, 2021).

R2 activity was also measured by changing growth media, and found to exhibit good regulatory performance in all tested media except SOC (Fig 5D). Less reduction of TurboRFP was detected when cultured in SOC. By carefully examining the composition of the media, we speculated that glucose in the SOC might prevent its function. To test this hypothesis, we added 0.4% glucose to the LB medium and subsequently measured its fluorescence. As expected, R2 was inactive in this medium. These data suggested a certain correlation between glucose concentration and theophylline transport. Since little is known about theophylline transport in *E. coli*, our results provided some clues for this study. In conclusion, our rationally designed TC-OFF theophylline riboswitch could control gene expression under a variety of bacteria, different growth media and temperatures, making it a useful tool for repressing gene expression.

### Enhanced repression by the dual transcription-translation control system R2-RepA

Through the above experiments, we found that although R2 had superior regulatory effect in *E. coli* JM101, up to 77.4-fold, the repression in the widely used ‘wild-type’ MG1655 strain was not ideal, only 10-fold at the maximum level of repression of 24 hours. To further reduce the leaky expression in MG1655, we introduced a second gene repression element into the system: a protein degradation tag (RepA), which consists of 15 amino acids (NQSFISDILYADIES) that directs the target protein to the housekeeping ClpAP protease (Butz *et al*, 2011; Hoskins *et al*, 2000). This element can be used to shorten the half-life of proteins, thereby reducing leaky expression. We considered it to be a promising regulatory tool because it is located at the N-terminus of the protein, so the RepA tag coding sequence could be easily integrated with the riboswitch coding sequence as a single regulatory cassette. We anticipated that the fusion protein called RepA-RFP could serve as a substrate for ClpAP degradation (Fig 6A), with the detailed sequences listed in Appendix Fig S3. The repression ability of the RepA tag was examined from 2 to 12 hours. As shown in Fig 6B, TuborRFP fluorescence intensities were indeed reduced in RepA-RFP (RepA, MG1655-pWA144) compared to untagged TurboRFP (no riboswitch, MG1655-pWA143), but leaky expression was still seen throughout growth. The lowest expression of RepA-RFP was at 6 hours, and the fluorescence gradually accumulated from 8 hours to 12 hours. One possible reason is that at stationary phase, the amount of proteins targeted for degradation by proteases increased, and they competed for a limited number of proteases, leading to a prolonged half-life of RepA-RFP (Zhou & Gottesman, 1998). We also tested the repression of TurboRFP by R2 from 2 to 12 hours. In the absence of theophylline, TurboRFP fluorescence intensities increased over time. After the addition of theophylline, they decreased from 1.4-fold at 4 hours to 6.5-fold at 12 hours (R2, MG1655-pWA140) (Fig 6C). However, the results showed that there was still significant leaky expression from 6 to 12 hours. Thus, we could see that neither RepA nor riboswitch alone could effectively inhibit gene expression.

**Figure 6.**
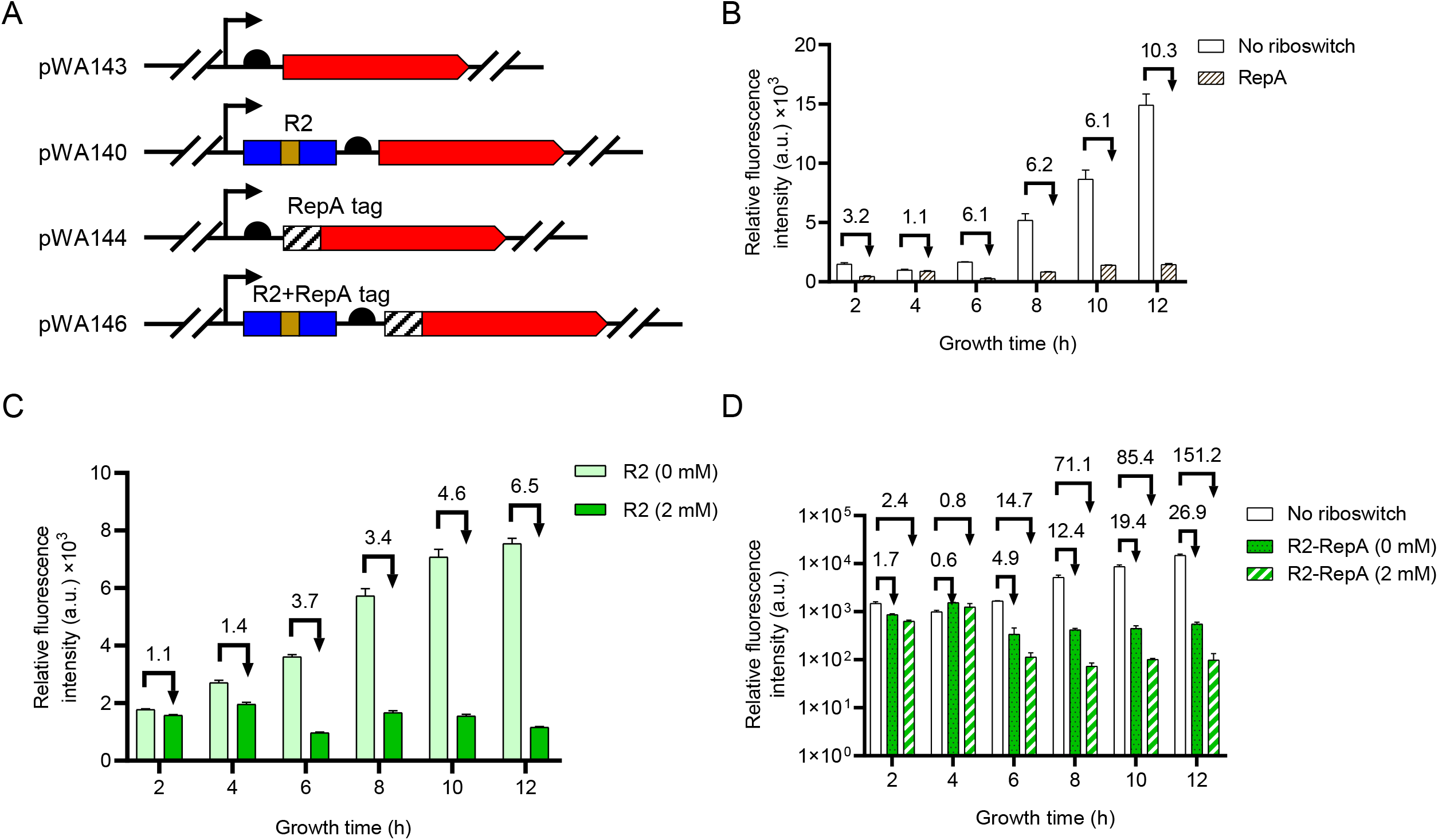
Regulation of TurboRFP expression by the R2-RepA system. A Schematic of plasmids containing the theophylline riboswitch coding sequences or RepA degradation tag coding sequences upstream of *turborfp*. pWA143 was used as a control plasmid without regulatory sequences upstream of *turborfp.* pWA140, pWA144 and pWA146 represent plasmids containing tandem riboswitch coding sequences, RepA tag coding sequences, and plasmids containing these two sequences, respectively. B Regulation of TurboRFP expression by protein degradation tag-RepA at different growth phases. The numbers above the column represent the TurboRFP fluorescence intensity of “no riboswitch” divided by “RepA” TurboRFP fluorescence intensity. C Regulation of TurboRFP expression by R2 at 0 or 2.0 mM theophylline over 2 to 12 hours. The numbers above the column represent the TurboRFP fluorescence intensity at 0 mM theophylline divided by TurboRFP fluorescence intensity at 2 mM theophylline. D Regulation of TurboRFP expression by R2-RepA system at 0 or 2.0 mM theophylline over 2 to 12 hours. The numbers above the column represent the TurboRFP fluorescence intensity of “no riboswitch” divided by TurboRFP fluorescence intensity at 0 mM or 2 mM theophylline, respectively. All data above represent mean ± SD of 3 biological replicates.

Next, the R2-RepA system, in which two theophylline riboswitch coding sequences in tandem were fused to the RepA coding sequence, was inserted upstream of *turborfp* (pWA146) (Fig 6A and Appendix Fig S3). We then measured the regulatory effect of R2-RepA system in media with or without theophylline. The strain produced nearly no fluorescence in the presence of theophylline (99 a. u.), highlighting the high repression efficiency of R2-RepA system (Fig 6D). As with R2-RepA system (R2-RepA, MG1655-pWA146), the maximum level of repression at 12 hours after addition of theophylline reached 151.2-fold compared to the system without the repression (no riboswitch, MG1655-pWA143). In another word, at 12 hours after adding theophylline, basal expression was particularly low, only 0.6% of TurboRFP alone. Our results supported the idea that the R2-RepA system is very effective in repressing gene expression.

## DISCUSSION

### A highly efficient dual gene expression regulatory system

We have successfully generated a new theophylline riboswitch that repress transcription on binding of theophylline. By combining transcriptional and translational regulation, we constructed a dual gene expression regulatory system based on riboswitches and protein degradation tag RepA, namely the R2-RepA system. The R2-RepA system is only 218 bp in length, and did not require additional expression of other regulatory proteins. The expression of any protein could now be repressed efficiently by simply inserting this new cassette upstream of the protein-coding sequence, followed by adding theophylline to achieve over 150-fold of gene repression. In addition, we systematically evaluated 25 theophylline riboswitches used in bacteria. We determined their activation/repression ratios and basal expression levels in various strains, growth media and temperatures. Therefore, these works provide the basis for a more rational selection of theophylline riboswitches.

### Possible reasons for the poor performances of theophylline riboswitches

In Appendix Table S4, we described in more detail than Table 1 all theophylline riboswitches developed in bacteria to date. We compiled information on sequences, lengths, regulatory mechanisms, growth conditions, construction methods, and activation/repression ratios, and tried to identify the causes of those riboswitches with poor performance (activation/repression ratio less than 2-fold), such as riboswitches No. 1, No. 3, No. 4, No. 15, No. 16, No. 17, No. 18 and No. 21. We first evaluated their full-length riboswitch structures by RNAfold (Gruber *et al*, 2008), and found that, of these riboswitches, the secondary structures of the two TL-ON riboswitches No. 1 and No. 21 were the most unstable, with a minimum free energy (MFE) of only −0.13 and −0.11 kcal/mol/bp, respectively. This means that, in the absence of theophylline, the RBS and start codon tended to be accessible for translation initiation, resulting in significant leaky expression. We also compared the structural differences between the full-length riboswitches and those with aptamer region only that were constrained to form efficient ligand-binding folds. TL-OFF riboswitch No. 3, RZ-ON riboswitch No. 4, TC-OFF riboswitch No. 15, No. 16 and No. 17 showed no secondary structure changes in either state. Therefore, we speculated that theophylline has little effect on maintaining the theophylline-binding secondary structure of these RNAs, resulting in no regulatory effect. We also calculated the free energy differences between theophylline-bound and theophylline-unbound state of these riboswitches. The free energy difference for No. 18 was −22.6, deviated significantly from the binding energy of the aptamer/theophylline complex (−8.86 kcal/mol) (Wachsmuth *et al*, 2013). Thus, this RNA did not appear to fold as the authors claimed.

### Applications of the dual transcription-translation control system R2-RepA

Compared to the numerous strategies to achieve high gene expression levels in *E. coli*, relatively few repression systems were available (Bervoets & Charlier, 2019; Kato, 2020). To date, three systems are commonly used in bacterial cells; the tetracycline-repression system (Tet-off system), clustered regularly interspaced short palindromic repeats interference (CRISPRi), and sRNA meditated gene repression. While useful in a large number of applications, these systems have limitations. The Tet-off system and the CRISPRi system require additional expression of the regulatory proteins TetR and Cas, which increase the manipulation difficulty and metabolic burden for bacteria (Hillen & Berens, 1994; Qi *et al*, 2013). Compared with these three repression systems, the current R2-RepA system had several advantages: 1) No additional protein expression is required; 2) The R2-RepA system is very short, only 218 bp in length; 3) There is less crosstalk between inducer and cellular metabolism (Yu *et al*, 2009); 4) The system is very effective, with up to 150-fold repression. Therefore, our system is simple, accurate and can be used as a general approach for repressing gene expression.

However, the R2-RepA system also had certain limitations. Since it is actually consisted of two different elements, the riboswitch and the protein degradation tag, we did not test the robustness of the protein degradation tag alone under various conditions. We hypothesized that proteases in different bacteria recognized different protein degradation tags, and therefore protein degradation tag was not universal in a wide range of bacteria. Actually, back in 2014, Cameron et al. developed a degradation system based on the *Mesoplasma florum* tmRNA system that can function in a wide range of bacteria. However, this system requires additional expression of the exogenous protease *mf*-Lon, which would increase the complexity of the system, so we did not adopt this system in this study (Cameron & Collins, 2014). We believe that any protein degradation tag corresponding to host bacterial protease can be used in conjunction with the theophylline riboswitch to repress gene expression, if desired.

### Future directions

To date, libraries of genetic regulatory elements with different regulatory strengths, such as promoters, RBS elements, and intrinsic terminators, have been constructed (Chen *et al*, 2013; Mutalik *et al*, 2013; Zaslaver *et al*, 2006). However, a library of theophylline riboswitches has not been established. So far, all theophylline riboswitches are derived from the same aptamer, and the modification of them are limited to the expression platform. Therefore, the screening of the theophylline riboswitch aptamer is also a very important aspect, and with the development of new technology after the systematic evolution of ligands by exponential enrichment (SELEX), we believe that more theophylline aptamers can be screened by new methods to obtain theophylline riboswitch with better performance. Meanwhile, studying the transportation mechanism of theophylline and improve its transportation efficiency can significantly improve its regulatory effect. This study thus represented a crucial step toward harnessing theophylline riboswitches and expanding the synthetic biology toolbox.

## APPENDIX DATA

Appendix Data are available at MSB online.

## ACKNOWLEDGEMENTS

This work was supported by the National Natural Science Foundation of China (31971339 and 32171422). And was also supported by the Fundamental Research Funds for the Central Universities (2662022SKYJ004).

